# A pipeline to track unlabeled cells in wide migration chambers using pseudofluorescence

**DOI:** 10.1101/2022.01.26.476896

**Authors:** Antonello Paola, Marcus Thelen, Rolf Krause, Pizzagalli Diego Ulisse

## Abstract

Cell migration is a pivotal biological process, whose dysregulation is found in many diseases including inflammation and cancer. Advances in microscopy technologies allow now to study cell migration *in vitro*, within microenvironments that resemble *in vivo* conditions. However, when cells are observed within large 3D migration chambers at low magnification and for extended periods of time, data analysis becomes difficult. Indeed, cell detection and tracking are hampered due to the large pixel size, the possible low signal-to-noise ratio and distortions in the cell shape due to changes in the z-axis position. Although fluorescent staining can be used to facilitate cell detection, it may alter cell behavior and suffer from fluorescence loss over time (photobleaching).

Here we describe the application of an image analysis pipeline based on deep learning to convert the transmitted light signal from unlabeled lymphoma cells to pseudofluorescence. Such pipeline confers a significant improvement in tracking accuracy while not suffering from photobleaching. This is reflected in the possibility of tracking cells for three-fold longer periods of time.

## Introduction

The regulation of many biological processes is mediated by the migration of cells from one anatomical location to another to exert their function. For example, primordial germ cell migration in zebrafish is essential to ensure the correct organ development ^1^. Moreover, the correct development of proper immune responses requires a fine-tuned regulation of leukocyte trafficking and migration^2–4^.

Amongst the mechanisms involved in cell migration, chemotaxis polarizes cells and controls the direction of migration toward favorable locations^5^, ^6^. Hence, several studies focus on the knowledge of the molecular mechanisms and signaling pathways that regulate chemotaxis *in vitro* and *in vivo*. However, the directional movement of cells is regulated not only by the type of soluble cues diffused into and retained by the environment, but also by the environment itself ^7, 8^.

Therefore, engineered microenvironments are essential to study cell migration *in vitro*. Amongst these, 3D migration is a setting where cells are embedded in collagen-like fibers to mimic the ECM matrix *in vitro*^6, 9^ (Fig. 1A). Widefield microscopy (WM) is an established technique to perform long-term imaging studies in wide migration chambers. WM can be performed by recording the intensity of the light transmitted through the sample, without necessarily requiring fluorescent staining. In this acquisition mode, the acquired data consists of a series of 2d grayscale images captured over time.

**Figure 1.**
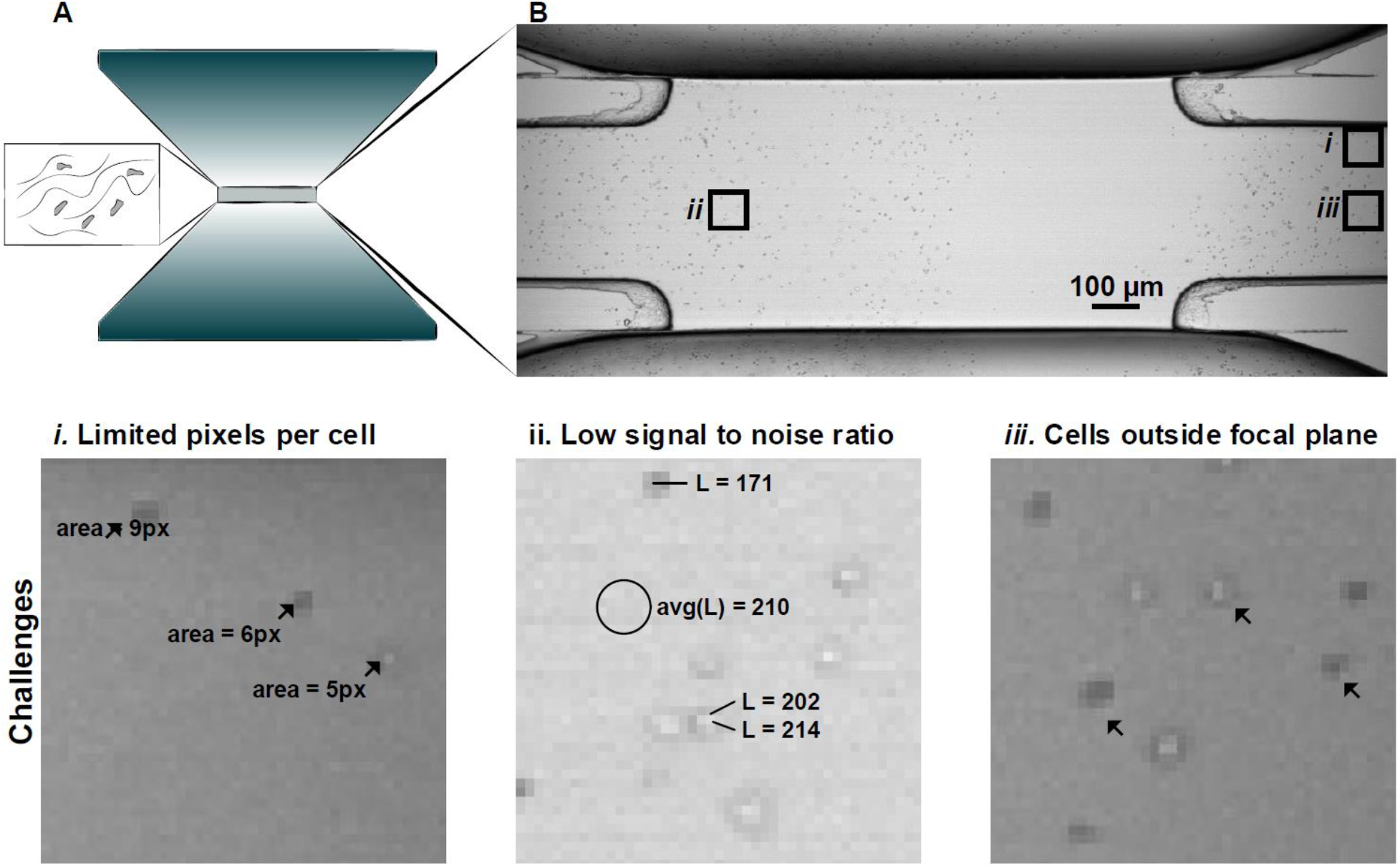
Widefield microscopy in wide 3D migration chambers. **A.** Representation of a wide migration chamber for chemotaxis assays. **B.** Widefield image of VAL cells scattered in the field of view using transmitted light. ***i-iii*** challenges for the automatic analysis: (*i*) large pixel size associated with a limited number of pixels per cell, (*ii*) appearance of cells similar to the background associated with a low signal to noise ratio, and (*iii*) different appearance of cells according to the position along the z-axis with respect to the focal plane.

However, when WM is applied to study the migration of highly motile cells such as leukocytes or metastasizing lymphoma cell lines, the analysis of the acquired series of images presents specific challenges. Indeed, the classical analysis pipeline involves three steps: cell detection, cell tracking, and computation of motility measures^10, 11^. The application of such a pipeline is hampered at the first step, due to dramatic changes in cell shape that introduce cell detection errors. These changes are associated with the frequent squeezing of the cytoskeleton during amoeboid migration through dense ECM ^6^, or introduced as an artifact during the migration along the z-axis. In the last case, cells are imaged outside the focal plane, leading to blurred and enlarged shapes in the acquired images.

Additionally, depending on the experimental settings, cells can require a long period to exert a directional movement. Hence, long acquisition times are needed. Long-time acquisitions may prevent the usage of fluorescent staining (used to facilitate cell detection) due to photobleaching. Hence, imaging of unlabeled cells using transmitted light (TL) is necessary. Lastly, analysis of cells following long tracks (i.e. > 150 μm), demands a large field of acquisition and necessitates low magnification (i.e. 4X objective). Therefore, the resolution is another problem that prevents efficient cell detection and subsequently compromises tracking (Fig. 1B).

Recent advances in artificial intelligence methods applied to bioimage analysis remarkably improved the accuracy of cell detection and tracking ^12, 13^. Although some of these methods can generalize to many different imaging modalities and cell types ^14^, a custom pipeline to automatize the analysis of image series capturing leukocytes using transmitted light with low resolution and in wide migration chambers is still missing.

Therefore, we propose WID-U (wide U-NET), a plugin for common bioimaging software such as Imaris (Oxford instruments) and FIJI, that converts the TL signal from widefield microscopy into pseudofluorescence. The pseudofluorescence signal generated by WID-U yielded an efficient detection of the cells using state of the art spot-detection and tracking algorithms such as TrackMate^15, 16^, and improved tracking accuracy of cells in challenging 3D *in vitro* environments.

## Results

### Pipeline to convert transmitted light to pseudo-fluorescence

To convert the transmitted light (TL) signal from unlabeled cells to pseudo-fluorescence, we developed an image processing pipeline based on deep learning. Such a pipeline was specifically developed to face the challenges arising when images of cells are acquired in large 3D migration chambers, at low magnification (4x) and large fields of view (2mm x 2mm). To account for such low magnification and large fields of view, images were processed with a sliding window of 56×56 pixels (~34μm x 34μm) (Figure 2, A, red square). To enable the subsequent application of deep learning, each window is interpolated and upscaled by a factor of 4 to 224×224 pixels, and processed via a patch classifier based on the U-NET architecture^17^ (Figure 2, B). Such architecture receives as input the upscaled TL images (Figure 2, B, grayscale image), and generates as output an image where the intensity of each pixel is the class one probability, or pseudofluorescence (Figure 2, B, magenta-colored image). The output of the U-NET is then downscaled and combined as a new imaging channel in the original image (Figure 2, C). A dataset consisting of 159 upscaled image pairs, with transmitted light and manually annotated binary masks of cells was created to train the U-NET (Figure 2, D).

**Figure 2.**
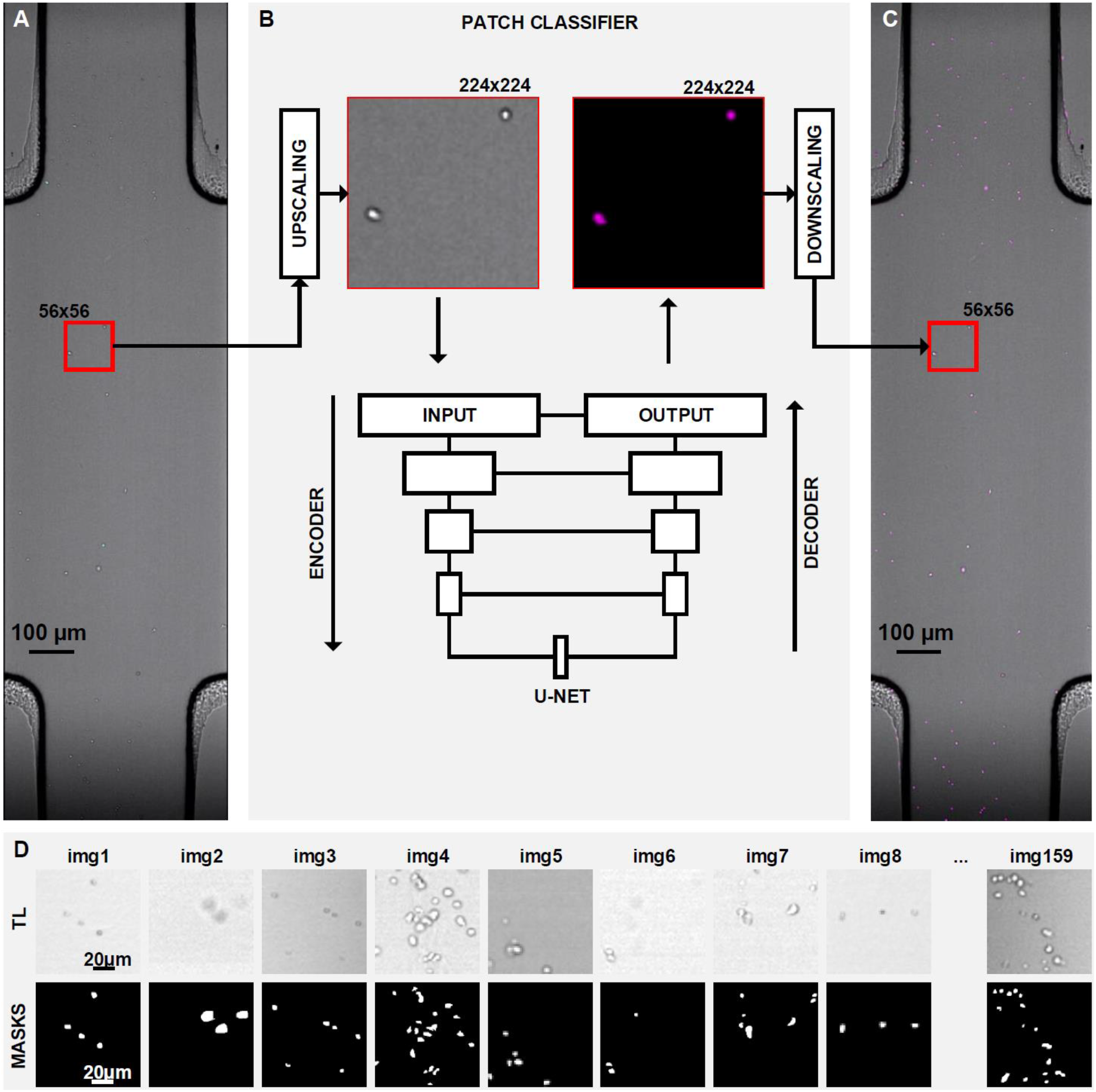
Pipeline to convert transmitted light to pseudofluorescence. **A.** TL image obtained by widefield microscopy. The red square represents a sliding window of 56×56 pixels used for image processing. **B.** Patch classifier, which employs a U-NET architecture with five fully convolutional layers. An upscaled window (224px x 224px) is used as input. The class-one probability is used to generate a pseudofluorescent image as output (magenta colored). **C.** The output is then downscaled and combined with the original imaging data to create a virtual imaging channel with pseudo-fluorescence (magenta). **D.** Representative image pairs from the dataset used for training, including transmitted light images (up), and manually annotated binary masks (bottom).(n of images included = 159).

### Enhanced cell detection and tracking using pseudo-fluorescence

We applied the proposed pipeline to analyze videos of VAL cells (a B cell lymphoma cell line), acquired in 3D microenvironments, and compared the quality of the pseudofluorescence signal with respect to transmitted light, or real fluorescence emitted by cyan fluorescent protein positive (CFP+) cells. The proposed pipeline yielded a significant improvement in the signal-to-noise ratio (SNR) with respect to TL images (Figure 3, A). Although SNR was higher, it was not significantly increased with respect to real fluorescence (CFP). However, in contrast to real fluorescence, pseudofluorescence did not suffer from photobleaching (Figure 3, B).

**Figure 3.**
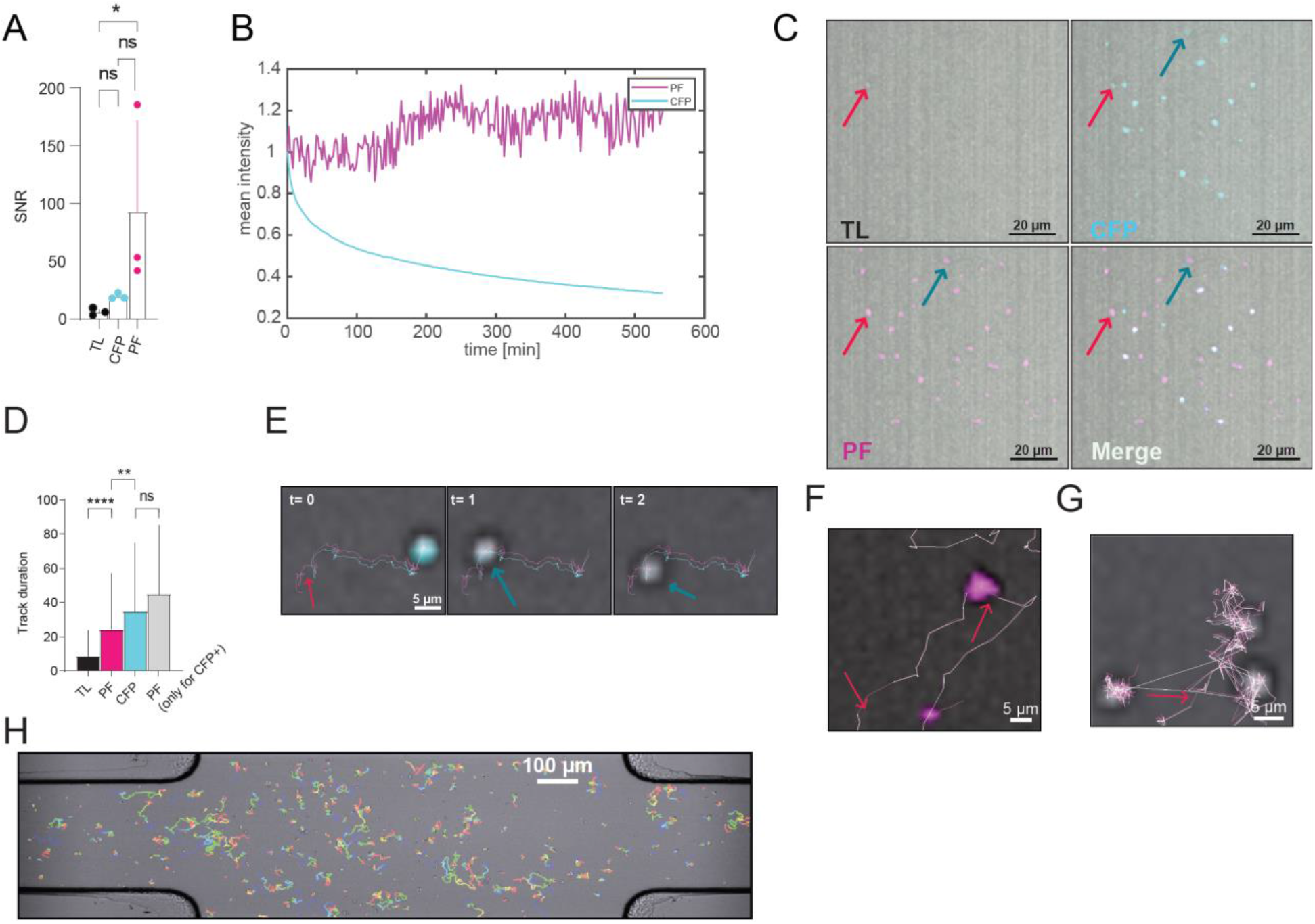
Enhanced cell tracking using pseudofluorescence. **A.** Comparison of the signal to noise ratio (SNR) between cells observed via transmitted light (TL), CFP labeled cells (CFP), and pseudofluorescence (PF). The SNR value of each population derives from the mean of three independent observations. **B**. Comparison of the fluorescence intensity over time of pseudofluorescence (magenta) with respect to CFP (blue), showing the effect of photobleaching on CFP. **C.** Representative micrographs showing the transformation into pseudofluorescence of cells which are poorly visible in TL and outside the focal plane. **D.** Comparison of automatic tracking accuracy (Track duration) using transmitted light, pseudo-fluorescence, or CFP-labeled cells. Values of each population were calculated from the mean of three independent observations. **E.** Representative micrograph showing tracks of cells obtained using PF (magenta lines) and CFP signals (cyan lines). **F.** Representative micrographs comparing tracking errors (red arrows, cells not detected) when using PF with respect to TL. Magenta-colored tracks were obtained with PF, white-colored tracks were obtained with TL. **G.** Representative micrograph showing tracking errors (red arrow, glitch between cells in close proximity) when using PF with respect to TL. **H.** Results obtained on a 3D migration chamber (color-coded tracks, blue = 0s, red = 8 hours). Statistic performed with ONE WAY ANOVA, *=P < 0.05 **=P<0.01 ****=P< 0.0001

Moreover, the proposed pipeline increased the visibility of cells, which were out of focus or with deformed shapes (Figure 3, C). Altogether, these properties make pseudofluorescence similar to real fluorescence, but with increased stability over time.

To validate the effect of pseudofluorescence on the quality of cell tracking we performed automatic cell tracking using transmitted light, real fluorescence (CFP+ cells), or pseudofluorescence signals. Pseudofluorescence yielded significantly more accurate tracks than the original transmitted light signal, with an average three-fold increase in the track duration (Figure 3, D). In comparison with real fluorescence, track duration was longer although not significant. More accurate tracks were obtained at later time points when the fluorescent signal was fading (Figure 3, B-E). In general, pseudofluorescence decreased the number of tracking errors, resulting in fewer interrupted tracks (Figure 3, F) and fewer glitches when cells were in close proximity (Figure 3, G).

Taken together, the novel method enabled the automatic tracking of cells using pseudofluorescence in large 3D microenvironments (Figure 3, H, Suppl. Figure 1).

## Materials and methods

### Cell line and cultures

VAL cells were cultured in medium supplemented with 10% heat-inactivated Fetal Bovine Serum (FBS), 1% Penicillin/Streptomycin, 1% GlutaMAX. 1% NEAA, 1% Sodium-Pyruvate, and 50 μM β-mercaptoethanol. CFP+ cells were cloned as described previously^18^.

### 3D migration

Migration assays of VAL cells were performed using the 3D chamber μ-Slide from Ibidi as described in *Antonello* et al. (submitted). Briefly, cells were embedded in a collagen matrix formed by 1.6 mg/mL PureCol (Collagen, Sigma-Aldrich), 0.36% PBS supplemented with 0.36% FBS, 0.036% P/S, 1.5 μg/mL recombinant human ICAM-1/CD54 Fc chimera (R&D systems) at 4° C. The temperature was slowly raised over 45 min to 37°C to induce a homogeneous collagen polymerization. Complete medium was added to both side reservoirs. After 24 hours 15 μL of 400 nM CXCL12 were added to one of the reservoirs and time-lapse video microscopy was performed for 6 hours at 20 seconds time intervals using an ImageXpress^®^ (Molecular Devices) high throughput microscope.

### Transformation of transmitted light images into pseudofluorescence

Transmitted light images acquired with a 4x objective were converted into pseudo-fluorescence by employing an end-to-end neural network with convolutional layers based on the U-NET architecture^17^. A dataset with pairs of transmitted light and pseudofluorescence (binary masks) was created by manually drawing the contours of cells. Such dataset included 150 images of 56×56 pixels (91×91 μm) from different experiments. These images were upscaled to 224 x 224 pixels to facilitate annotation then downscaled to 112×112 pixels for training and augmented to 15’000 images. The trained network was then applied to convert images of size >= 1000×500 pixels, to pseudofluorescence by classifying a moving window of 56×56 pixels. The class-1 probability computed by the U-NET was used as pseudofluorescence.

### Cell tracking

Initially, TL images were transformed to pseudofluorescence images. Then, cells were detected and tracked using the Spots tracking functionality of the Imaris software (Oxford Instruments, v.9.7.2) in the original TL channel, in the imaging channel capturing fluorescence, and in the pseudofluorescence channel. In all cases, an estimated spot diameter of 8 um was selected and background subtraction was enabled to account for non-uniform illumination. Tracking was performed using an autoregressive motion model, with a maximum distance of 20 μm, and a maximum gap size of 0. Tracks shorter than 300 seconds were excluded from the analysis. Tracks were divided into two classes (WT and CFP+ cells), based on the mean fluorescence intensity of the imaging channel centered on CFP. Finally, the duration of each track was computed. Tracks outside the migration channel were deleted manually.

Automatic tracking was also performed using TrackMate in FIJI^16^ using the LoG spot detector or automatic thresholding, and Simple Lap tracker for spot tracking. The same values described before for the tracking in Imaris were used also for the tracking in TrackMate.

### Data and code availability

The software to convert transmitted light into pseudofluorescence is available in Supplementary Files, u4x.zip, including the executables and the source code in Matlab, Batch, Bash, and Python. The package includes a plugin for the imaging software Imaris (Oxford Instruments), a trained U-Net model, and the scripts to train a custom model. The U-NET model was defined and trained in the Keras (v 2.3.1) framework, on a TensorFlow-GPU (v 2.2.0) backend, using Python (v 3.8.3). A CUDA-enabled machine is recommended for the execution of the network. To configure the connection to such a machine, instructions are included in the README file. The program to create the training dataset with upscaling and manually drawing of binary masks was written in Matlab r2019b, and is available in the file “annotate_dataset.m”. The dataset used to train the U-NET architecture is available in training_data.zip. Data and code is released under the Open Source GPL v3 license at https://github.com/pizzagalli-du/wid-u

### WID-U usage

To facilitate the execution of the pipeline for transmitted light to fluorescence conversion, a plugin for the Imaris (Oxford instruments), and FIJI bioimaging software has been developed. The usage of the plugin is summarized in (Figure 3) and requires a deep-learning enabled machine for image processing.

### Statistics

SNR and Track duration values were analyzed with PRISM software. Statistics performed with ONE WAY ANOVA * P<0.05, **P<0.01 ****P<0.0001.

## Discussion

The application of deep learning to *in vitro* time-lapse imaging improved the tracking accuracy of leukocytes in large 3D microenvironments. It was accomplished by training a U-NET architecture with a custom dataset that included B lymphocytes. To apply our pipeline to other cell types or other imaging modalities, the dataset can be extended including images acquired with those settings and the network re-trained. To minimize the number of training examples required to adapt the network to other conditions we recommend the usage of transfer learning approaches^19^. The creation of the training set was facilitated by upscaling. This is associated with an increased precision during the manual annotation when cells had a small area (i.e. 4 pixels).

The tracking was improved as a consequence of more accurate spot detection. The proposed pseudofluorescence transformation can be combined with a variety of tools for spot detection. In this work, we tested two approaches typically used in bioimaging: watershed with background subtraction (Imaris) and LoG detector (FIJI/TrackMate). In both cases, performances improved. Performances increased also when automatic thresholding (FIJI/TrackMate) was used for spot detection. This suggests that pseudofluorescence is uniform across the field of view and detection of objects based on intensity values rather than morphological features is possible. To further enhance accuracy, pseudofluorescence can be used in combination with recently developed methods for cell detection, such as those based on geometric properties and deep learning^20^.

In conclusion, the proposed pipeline allowed cell tracking in large 3D migration chambers over extended periods of time, retaining the simplicity of cell detection as when real fluorescence is used, but avoiding photobleaching and other side effects of cell labeling.

---

**Supplementary Figure 1.**
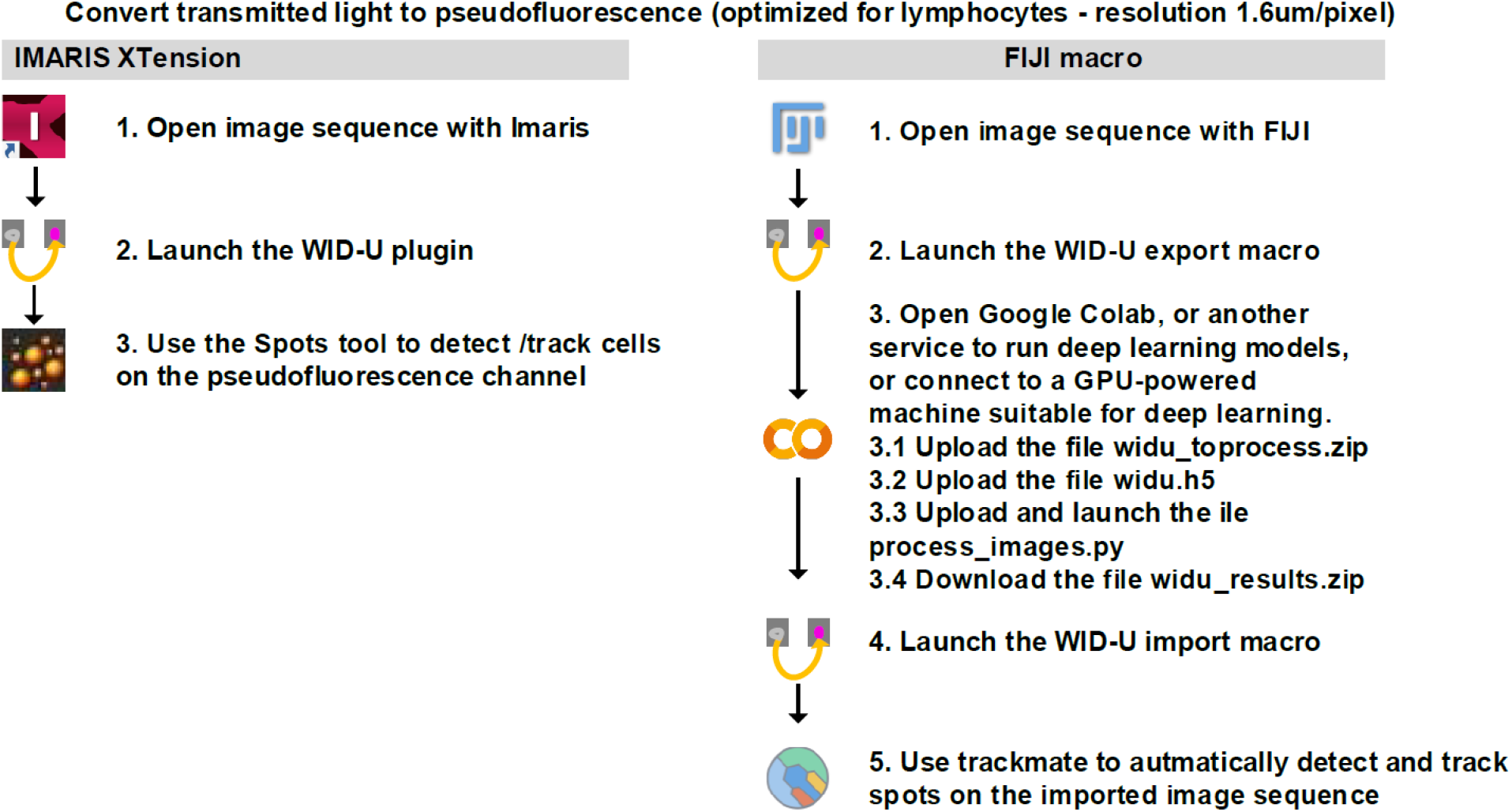
Usage workflow. The pipeline to convert low-resolution transmitted light images to pseudofluorescence is made available via the WID-U plugin for the bioimaging software Imaris (left) and FIJI. In Imaris, once installed and configured for communication with a deep learning-enabled machine, the user can launch the plugin and use the Spots tool for cell detection/tracking. In FIJI, once the image has been opened, the user can launch the WID-U export macro to export and pack the data to be processed. Data need to be uploaded to a deep learning-enabled machine along with the network weights (widu.h5) and Python code for image processing (process_images.py). Once completed, the file widu_results.zip is created and can be downloaded/imported in FIJI using the WID-U import macro. The TrackMate plugin can be used for spot detection and tracking on imported images.

**Supplementary Figure 2.**
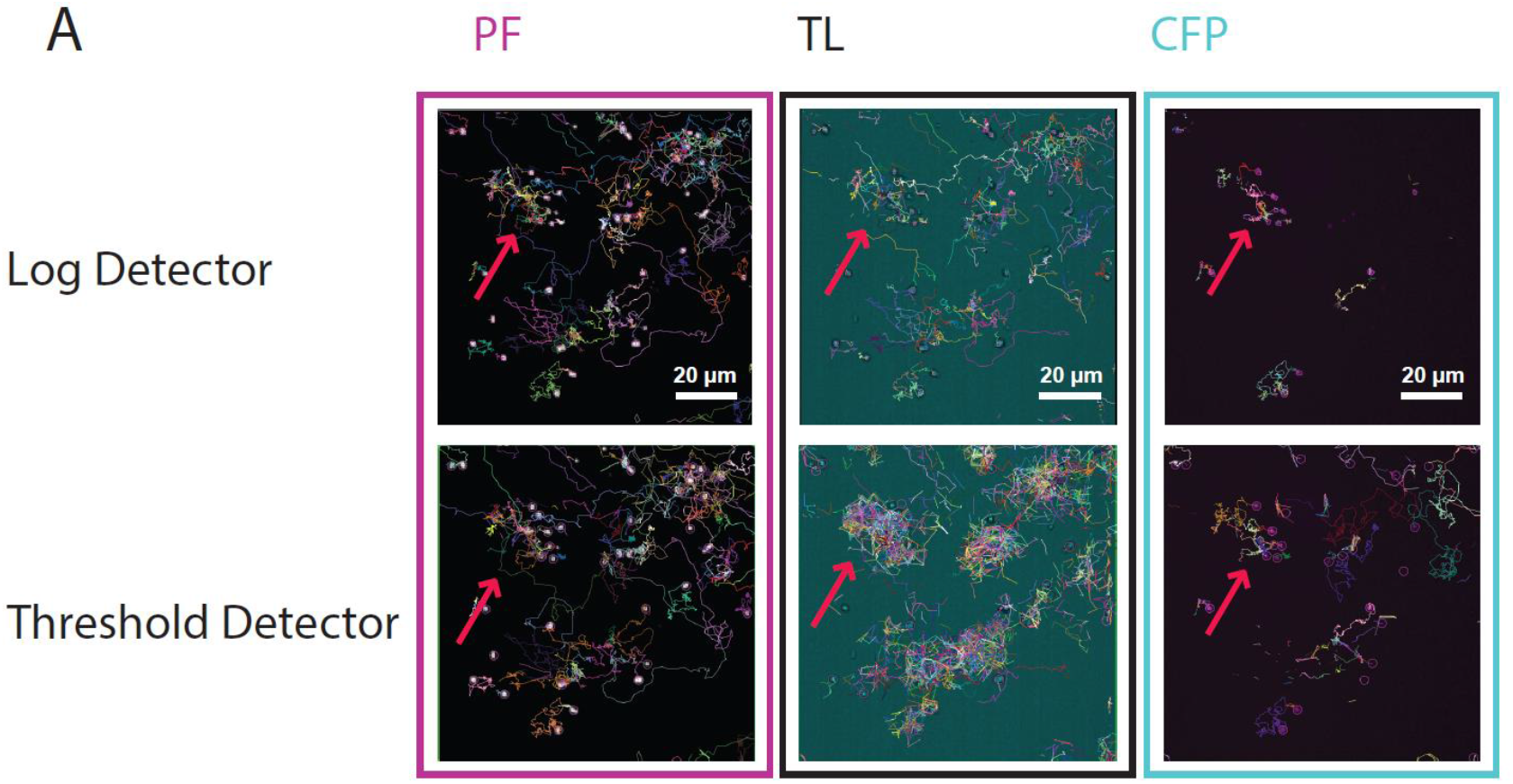
Representative micrograph showing tracks of cells obtained with PF (magenta square), TL (black square), or CFP (cyan square), using Trackmate analysis. Detection of cells was performed using either LogDetector (above) or Threshold detector (below).

## Notes

### Competing Interest Statement

The authors have declared no competing interest.

## References

1. Boldajipour, B. et al. Control of Chemokine-Guided Cell Migration by Ligand Sequestration. Cell 132, 463–473 (2008).

2. Luster, A.D., Alon, R., & Von Andrian, U.H. Immune cell migration in inflammation: present and future therapeutic targets. Nat. Immunol. 6, 1182–1190 (2005).

3. Madri, J.A. & Graesser, D. Cell migration in the immune system: the evolving inter-related roles of adhesion molecules and proteinases. Dev. Immunol. 7, 103–116 (2000).

4. Mayor, R. & Etienne-Manneville, S. The front and rear of collective cell migration. Nat. Rev. Mol. Cell Biol. 17, 97–109 (2016).

5. Baggiolini, M. Chemokines and leukocyte traffic. Nature 392, 565–568 (1998).

6. Yamada, K.M. & Sixt, M. Mechanisms of 3D cell migration. Nat. Rev. Mol. Cell Biol. 20, 738–752 (2019).

7. Sengupta, S., Parent, C.A., & Bear, J.E. The principles of directed cell migration. Nat. Rev. Mol. Cell Biol. 22, 529–547 (2021).

8. De la Fuente, I.M. & Lopez, J.I. Cell Motility and Cancer. Cancers. (Basel) 12, (2020).

9. Sixt, M. & Lammermann, T. In vitro analysis of chemotactic leukocyte migration in 3D environments. Methods Mol. Biol. 769, 149–165 (2011).

10. Pizzagalli, D.U. et al. Leukocyte Tracking Database, a collection of immune cell tracks from intravital 2-photon microscopy videos. Sci. Data 5, 180129 (2018).

11. Beltman, J.B., Maree, A.F., & de Boer, R.J. Analysing immune cell migration. Nat. Rev. Immunol. 9, 789–798 (2009).

12. Ulman, V. et al. An objective comparison of cell-tracking algorithms. Nat. Methods(2017).

13. Berg, S. et al. ilastik: interactive machine learning for (bio)image analysis. Nat. Methods 16, 1226–1232 (2019).

14. Stringer, C., Wang, T., Michaelos, M., & Pachitariu, M. Cellpose: a generalist algorithm for cellular segmentation. Nat. Methods 18, 100–106 (2021).

15. Ershov, D. et al. Bringing TrackMate into the era of machine-learning and deep-learning. bioRxiv2021 (2021).

16. Tinevez, J.Y. et al. TrackMate: An open and extensible platform for single-particle tracking. Methods 115, 80–90 (2017).

17. Ronneberger, O., Fischer, P., & Brox, T. U-net: Convolutional networks for biomedical image segmentation. International Conference on Medical image computing and computer-assisted intervention, 234–241. 2015. Springer. Ref Type: Conference Proceeding

18. Puddinu, V. et al. ACKR3 expression on diffuse large B cell lymphoma is required for tumor spreading and tissue infiltration. Oncotarget. 8, 85068–85084 (2017).

19. Falk, T. et al. U-Net: deep learning for cell counting, detection, and morphometry. Nat. Methods 16, 67–70 (2019).

20. Fazeli, E. et al. Automated cell tracking using StarDist and TrackMate. F1000Res. 9, 1279 (2020).

